# Digestate as Biofertilizer for the Growth of Selected Vegetables and Illumina analysis of Associated Bacterial Community

**DOI:** 10.1101/2020.12.03.393058

**Authors:** F.C. Raymond, O.M. Buraimoh, O.S. Akerele, M.O. Ilori, O.T. Ogundipe

**Affiliations:** Department of Microbiology, Faculty of Science, University of Lagos, Akoka, Lagos, Nigeria; Department of Botany, Faculty of Science, University of Lagos, Akoka, Lagos, Nigeria

**Keywords:** Biofertilizer, Organic biomass, Digestate, Illumina sequencing, Nitrogen fixer

## Abstract

Chemical content of crops above desirable level, high cost, in addition to land and water pollution is a major drawback of applying chemical fertilizers. In this study, Digestate was used as a biofertilizer for the growth of selected vegetables. Furthermore, Illumina platform was employed to unravel the bacteria community of the digestate. *Corchorus olitorius*, *Amaranthus hybridus*, *Bot-celosia argentia* and *Talinum triangulare* were grown in 16 experimental pots for 35days using cured digestate. Soil treated with chemical fertilizer was used as a positive control while the soil without any fertilizer was used as a negative control. The plant height of *Talinum triangulare* for soil treated with digestate was 23.5cm and 34cm by days 7 and 35 respectively after plant emergence. *Corchorus olitorius* had stunted growth under same treatment. Whereas, *Bot-celosia argentia* and *Amaranthus hybridus* grew poorly on all treatments. The statistical analysis showed a significant difference (p≤0.05) between *Talinum triangulare* grown in soil treated with digestate (plant height = 23.50, plant number =10 at day 7) compared with those treated with NPK (plant height = 18.50, plant number = 6.50 at day 7). The plant height and number for untreated soil at day 7 were 10.90 and 5.0 respectively). The Illumina sequencing of the digestate revealed the presence of some beneficial soil bacteria including *Clostridium, Bacillus, Pseudomonas, Actinobacteria,* and *Micrococcus*. The presence of these bacteria known to be Nitrogen fixers and Phosphate solubilizers confer biofertilizer potential to the digestate.

## 1.0 INTRODUCTION

Bio-fertilizers are one of the best modern agricultural instruments and are seen as a gift from modern agricultural science to turn waste into functional types. Bio-fertilizers are environmental friendly fertilizers that not only avoid harm to natural sources, but often help clean up nature from precipitated chemical fertilizers to some degree. In the agricultural sector, these bio-fertilizers are used as a substitute for chemical fertilizers that are not eco-friendly and can destroy soil fertility in the long term (Owamah *et al*., 2014). Bio-fertilizers are preparations containing latent cells of effective microorganisms that assist the absorption of nutrients by crop plants when applied via seed or soil by their interactions in the rhizosphere (Di Maria *et al*., 2017; Yasar *et al*., 2017).

Digestate is composed of microbial biomass, semi-degraded organic matter and inorganic compounds and is derived from anaerobic digestion (AD) of organic waste. Digestate can be used as a biofertilizer for fertilization of farmlands, as an inorganic fertilizer or raw material for biofertilizer production. (Alburquerque *et al*., 2012; Roopnarain and Adeleke, 2017). AD is a regulated degradation of organic waste in the absence of oxygen and in the presence of anaerobic microorganisms (Ojolo *et al*., 2007). AD is also a means of dealing with organic waste and at the same time meeting global energy needs, it minimizes bulk of organic matter to be disposed, generates digestate which is rich in nutrients and has agricultural value and produces biogas rich in methane which can be used directly as fuel or converted to compressed natural gas and liquefied natural gas. Over dependence on inorganic chemical fertilizers has led to decline in soil quality, eutrophication and heavy metals pollution (Zhu *et al*., 2010; Owamah *et al*., 2014). Therefore, in the provision of environmental benefits like improvement of soil, food quality and protection as well as human and animal health, digestate as bio-fertilizers are necessary (Johansen *et al*., 2013). With several studies showing similar or higher yields in the use of digestate as an alternative to chemical fertilizer (Nkoa, 2014), this provides farmers with a new strategy that enables them to achieve the targeted objective of food security in Nigeria by growing food grain yields of high productivity (Food and Agricultural Organization, 2012). There are various types of digestate as bio-fertilizers and their key differences are generally in the raw materials used for their processing, forms of utilization and the source of microorganisms used in the preparation (Garfi *et al*., 2011). In addition, the consistency of the digestate can be determined by both the organic and inorganic matter present in the substrates. During the AD procedure, the microbial communities consume much of the organic matter and transform some to inorganic compounds. In the digester, for instance, the available nitrogen, whether from the substrate, atmosphere or from purging, is converted into ammonium and nitrates that remain in the digester until the completion of the AD process. The anaerobes in the digester do not use the inorganic matter during AD. Ammonium, an inorganic component which is essential for plant uptake is present in the digestate, thus makes digestate suitable for use as a fertilizer/soil conditioner (Bowen *et al*., 2014).

Digestate usually contains microorganisms like *Pseudomonas, Klebsiella, Salmonella, Penicillium, Shigella, Bacteroides, Aspergillus*, *and Bacillus* etc. some of which can be exploited in the production of bio-fertilizers because they hasten the microbial processes in the soil and increase the availability of nutrients that can be assimilated by plants (TNAU, 2008). *Klebsiella* and *Clostridium* species are free living nitrogen fixers while *Bacillus* and *Pseudomonas* species are phosphate solubilizers (Alfa *et al*., 2014). Digestate contains more readily available nutrients than non-digested products which make it better for crop fertilization (Garfi *et al*., 2011). In comparison to chemical fertilizers requiring high costs, various raw materials such as rural, municipal and domestic waste are sufficient for the inexpensive production of digestate as biofertilizers (Curry and Pillay, 2012; Dai *et al*., 2013). To this point, in many cropping systems around the world, the use of fiber and liquor from anaerobic digestion has contributed to increased fertilizer use and thus less chemical use (Sun *et al*., 2015). In the meantime, some studies have indicated that when used in conjunction with soil solarization, controlled management of digestate can minimize soil-borne disease and help in the suppression of weed seeds (Fernandez-Bayo *et al*., 2017).While anaerobic digestion has been shown to inactivate some soil pathogens, others, such as Listeria and some spore formers, have been documented to be less affected by anaerobic digestion (Insam *et al*., 2015). In so far as digestate is found to contain a high proportion of mineral nutrients, care must be taken to avoid over-use on land which may lead to environmental effects such as nutrient runoff, phytotoxicity, exposure to pathogens, accumulation of metals or increased exposure to NH_3_ gas (Nkoa, 2014; Insam *et al*., 2015). Treatment is needed to further eliminate pathogens and to generate qualitative digestate. Regulations and the expected end-use are the key factors for the treatment of digestate. Digestate can be used as fertilizer after removal from the digester without further treatment. However, in this case, the storage, transport, handling and use of digestate as fertilizer results in considerable costs for farmers compared to the value of the fertilizer due to its high volume and low dry matter. Digestate processing can be approached in two ways. The first is digestate conditioning, which aims to create a standardized biofertilizer (solid or liquid) that increases the consistency and marketability of the digestate. The second can be defined as digestate treatment similar to wastewater treatment; it is used to extract nutrients and organic matter from the effluent and to allow safe discharge. In most cases, both conditioning and treatment would be required in order to create a viable digestate process. (Drog *et al*., 2015).

## 2.0 Materials and Methods

### 2.1 Sample preparation and collection

The organic wastes (substrate) used for the anaerobic digestion include sawdust, fruit and food waste while cow dung was the source of microbial inoculum. Fresh cow dung was collected from Bariga market abattoir in Lagos state, Nigeria. Fruit waste (pineapple peel) and food waste (leftover rice) were collected from the fruit and food vendors within the University of Lagos premises. Sawdust was also collected from a timber shop at Bariga, Lagos. The anaerobic digester pilot setups were carried out at the Department of Microbiology, University of Lagos Nigeria. After 21 days of digestion and collection of methane, the end product (digestate) was collected into a sterile polythene bag and used as biofertilizer in pot trial experiments, while some samples of the digestate was collected into a sterile sample collection tube using a sterile spatula for nutritional analysis and metagenomic analysis using illumina platform.

### 2.2 Soil and digestate characterization

#### Analysis of major nutrient content of digestate and soil samples

Major elements including Nitrogen, phosphorus and potassium (NPK) composition of the digestate and soil samples used in this study was analyzed, sandy loam soil of low nutrient (<1% Nitrogen determined via analysis) was used. This soil type was chosen to ensure effective evaluation of the potency of the applied biofertilizers. The nutrient analysis of the soil and digestate was carried out at the Biochemistry Laboratory, University of Lagos teaching hospital (LUTH), Idi Araba, Lagos, Nigeria, before planting to determine the soil pH, Nitrogen (N), phosphorus (P) and Potassium (K) constituent. Total Nitrogen (N) was determined using Kjeldal method in a digestion and distillation method and the color derivative is measured using a spectrophotometer. The Phosphorus (P) was determined using Vanado Molybdate Method, where 5ml of the sample is diluted in 50ml volumetric flask and 10 ml of Vanado Molybdate is added with some water to make up the 50 ml mark and allowed to stand for about 10 minutes then the absorbance is read on a spectrophotometer. The Potassium (K) was measured using atomic absorbance spectrophotometry (AAS). The nutrient analysis was done to ensure accurate data after the experiment. The digestate was cured by storing for 21 days in sterile sacks in wet forms at room temperature prior to field application (Alfa *et al*., 2013) and the soil was sterilized using hot air oven at 118°C. Three (3) kg of soil was used per pot experiment and mixing of the digestate which was 1.5 kg with the soil was done and allowed to incubate for two weeks before commencement of planting. Commercially purchased chemical fertilizer (NPK 20-20-20) was used as positive control while an experiment without any fertilizer application was also set up as negative control as previously reported by Trinchera *et al*. (2013).

### 2.3 Pot trial experiment (the planting procedure)

3 kg of soil to 1.5 kg of the digestate was used for this experiment, this was placed in the experimental pot incubated for two weeks before planting. The experiment was carried out from the 8^th^ of May 2019 to 28^th^ June 2019. Four pots per plant, labelled; Soil + digestate (A), Soil + digestate (B), Soil + NPK (positive control), Soil only (negative control) were used, one of the plant pot contain the sterilized soil plus digestate (treated soil), another containing soil only as a negative control with no fertilizer (untreated soil), and a positive control using the chemical fertilizer. The set up can be seen in figure 1 below. The seeds of the vegetables were sown via spraying method on the soil. The ewedu (*Corchoris olitorius*) seeds were soaked in hot water for some minutes, air dried then planted using spraying method. These vegetables were grown in a controlled environment by putting certain factors into consideration, such as amount of water and exposure to direct sunlight. Water was sprayed on the plants morning and evening on daily basis and the plants were placed in improvised garden covered with a transparent polythene to minimize direct exposure to sunlight, as seen in figure 1 below. The data was collected every 7-days after seed emergence (DAE) on the following phyto-parameters: Leaf number per stem, total Leaf area, Plant height, and number of plant that emerged as previously described by Makadi *et al*. (2008).

### 2.4 DNA extraction and Illumina sequencing

Extraction of DNA was carried out at African Biosciences Ibadan, Oyo State Nigeria. Sequencing was carried out by a commercial laboratory (Xcelris Labs Limited Bodakdey, Ahmedabad Gujarat, India). The procedures include: DNA extraction, PCR amplification of the 16SrRNA, Gel electrophoresis and Sequence analysis. The DNA extraction was carried out initially, using Presto Soil DNA Extraction kit (Gene aid), which involves different extraction stages to obtain the purified DNA. The DNA was amplified as advised by Xcelris genomics lab, the amplification was made using 16SrRNA Primers, the PCR cycling condition was as follows: denaturation at 95°C for 3minutes, 35cycles of denaturation at 95°C for 30seconds, annealing at 54°C for 30 seconds and extension at 72°C for 30 seconds, with a final extension at 72°C for 10 minutes. The primer sequence used for the PCR is shown in table 2 below.

**Table 1:** treatments used in pot experiment.

| Codes | treatments |
| --- | --- |
| 1 | soil |
| 2 | soil + digestate |
| 3 | soil + chemical fertilizer (NPK) |

**Table 2.**
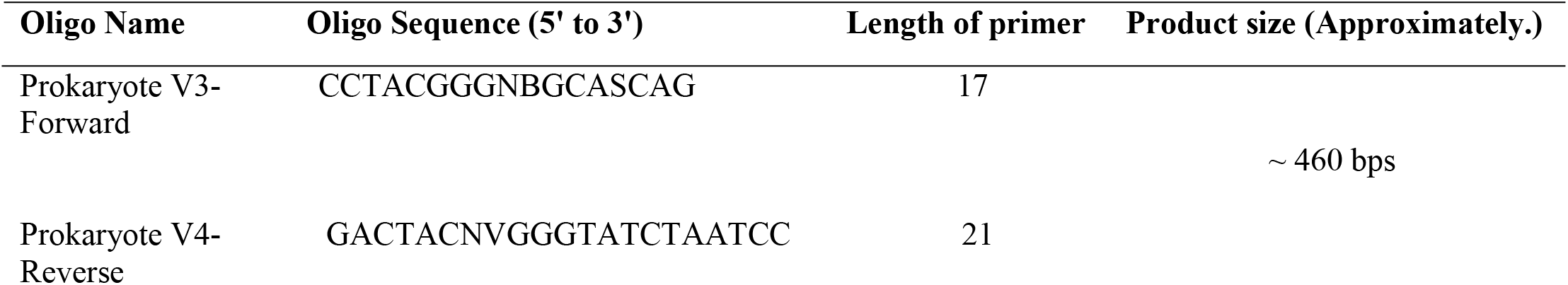
Primer sequence used for PCR.

The metagenomic sequencing was performed using Nextera XT Index Kit (Illumina Inc.) as well as the 16S Metagenomic Sequencing Library preparation protocol (Part # 15044223 Rev. B). Primers for the amplification of the V3-V4 hyper-variable region (Table 2) of 16SrRNA gene of bacteria were designed in house by Xcelris Labs Limited. These primers were synthesized in Xcelris PrimeX facility. The amplicon with the Illumina adaptors were amplified by using i5 and i7 primers that add multiplexing index sequences as well as common adapters required for cluster generation (P5 and P7) as per the standard Illumina protocol. The amplicon libraries were purified by 1X AMpureXP beads, checked on Agilent DNA1000 chip on Bioanalyzer2100 and quantified by Qubit Fluorometer 2.0 using Qubit dsDNA HS Assay kit (Life Technologies). After obtaining the Qubit concentration for the library and the mean peak size from Bioanalyzer profile, library was loaded onto Illumina platform at appropriate concentration (10-20pM) for cluster generation and sequencing. Paired-End sequencing allows the template fragments to be sequenced in both the forward and reverse directions on Illumina platform. The kit reagents were used in binding of samples to complementary adapter oligos on paired-end flow cell. The adapters were designed to allow selective cleavage of the forward strands after re-synthesis of the reverse strand during sequencing. The copied reverse strand was then used to sequence from the opposite end of the fragment.

### 2.5 Statistical analysis

The data of the different soil treatments for waterleaf plant (*Talinum triangulare*) were analyzed using GraphPad Prism 7.2, which allowed the simultaneous evaluation of the plant parameters between the different soil treatments.

## 3.0 RESULTS

### 3.1 Nutritional analysis of soil, digestate and chemical fertilizer

Table 3 shows the results of the nutrient analysis carried out on the soil and digestate prior planting indicating that the digestate is rich in Nitrogen having 1.60% as compared to the soil alone which had 0.94%, the mixture of the two (soil + digestate) increased the Nitrogen content of the soil (1.62%). This results shows a difference in pH and major nutrient components such as Nitrogen (N), Phosphorus (P), and Potassium (K) between the digestate, chemical fertilizer and soil. The digestate had a high pH value which happen to be more alkaline than the pH of the chemical fertilizer, the percentage of Nitrogen present in the digestate was higher than that of the soil only which made the mixture of soil and digestate to also have higher percentage of Nitrogen (N). The soil had higher concentration of other components such as potassium and phosphorus than in the digestate.

**Table 3:** Nutritional Content of the major plant nutrient in the digestate and chemical fertilizer.

| Experimental setups | pH | Nitrogen N (%) | Phosphate P (mg/100g) | Potassium K (ppm) |
| --- | --- | --- | --- | --- |
| Soil only | 6.0 | 0.94 | 50.65 | 67.41 |
| Soil + digestate | 7.9 | 1.62 | 16.11 | 42.67 |
| Digestate only | 8.1 | 1.60 | 18.44 | 55.31 |
| Chemical fertilizer (NPK) | 7.2 | 20 | 20 | 20 |

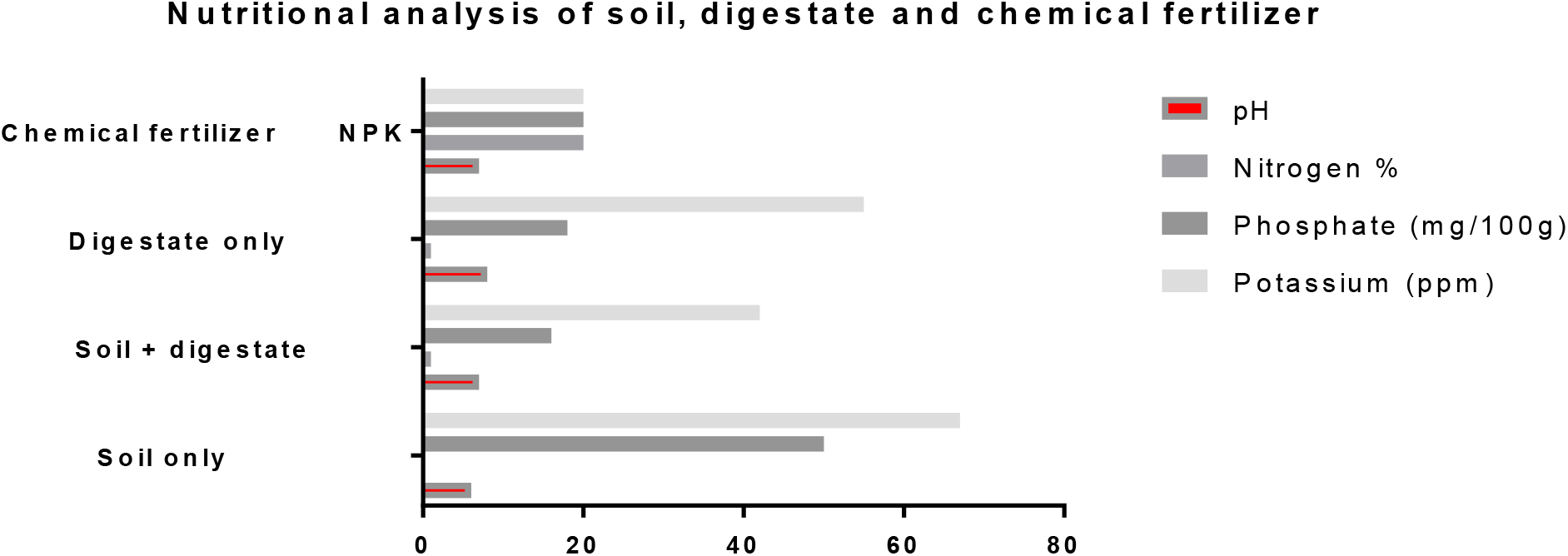

### 3.2 The vegetables yield

#### 3.2.1 Tabular presentation of Ewedu (*Corchorus olitorius*) growth parameter on different treatments

The growth parameter of (*Corchorus olitorius*) on different treatments were tabulated as seen in table 4 below. There was a significant increase in plant height and plant number on the treatments with the chemical fertilizer and on soil only as compared to that of the soil with digestate in the first 14 to 28 day after plant emergence (DAE). Figure 2 below shows the Ewedu plant at 35 days after plant emergence.

**Figure 2:**
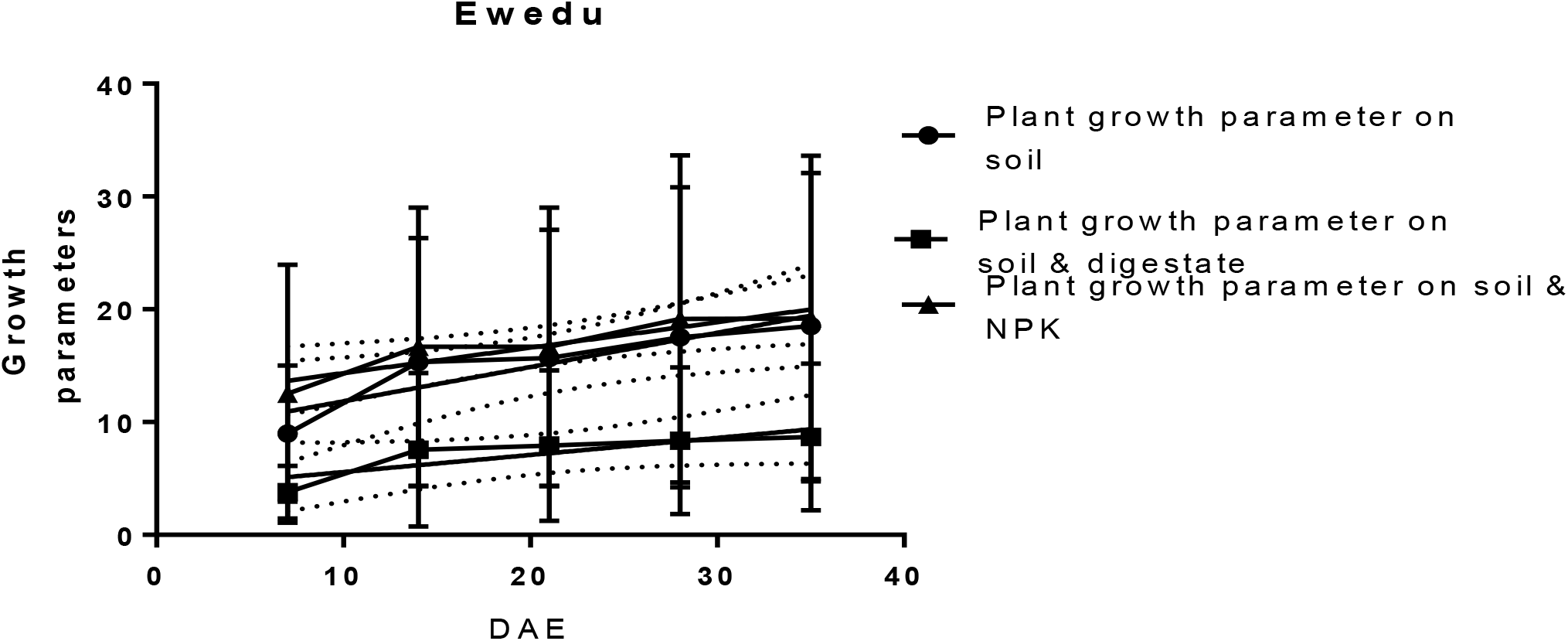
Ewedu (*Corchorus olitorius*) at 35 days

**Table 4:** Tabular presentation of the growth parameter for Ewedu (*Corchorus olitorius*) on different treatments from day 7 to 35 DAE.

| Days After Plant emergence (DAE) | Plant growth parameter on soil only |  |  |  | Plant growth parameter on soil & digestate |  |  |  | Plant growth parameter on soil & NPK |  |  |  |
| --- | --- | --- | --- | --- | --- | --- | --- | --- | --- | --- | --- | --- |
|  | No. of plant that emerged | Plant height (cm) | Total leaf area (cm<sup>2</sup>) | No. of leaves per stem | No. of plant that emerged | Plant height (cm) | Total leaf area (cm<sup>2</sup>) | No. of leaves per stem | No. of plant that emerged | Plant height (cm) | Total leaf area (cm<sup>2</sup>) | No. of leaves per stem |
| 7 | 9 | 15 | 2.9 | 2 | 4 | 6 | 1.3 | 2 | 10 | 25 | 2.5 | 2 |
| 14 | 19 | 24 | 2.9 | 5 | 15 | 6 | 1.6 | 4 | 20 | 27 | 3.0 | 5 |
| 21 | 19 | 25 | 3.0 | 5 | 15 | 7 | 1.7 | 4 | 20 | 27 | 3.0 | 5 |
| 28 | 19 | 30 | 3.5 | 8 | 15 | 8 | 2.0 | 4 | 22 | 32 | 3.4 | 8 |
| 35 | 22 | 30 | 3.5 | 8 | 15 | 9 | 2.0 | 4 | 22 | 32 | 3.5 | 10 |

#### 3.2.2 Tabular presentation of 12 on different treatments

Table 5 below reveals an interestingly increase in the plant grown on soil & digestate with an increase in plant height from 23.5cm from day 7 to 34cm at 35DAE as compared to the plant height on soil & NPK and soil only. Apparently the water leaf had a good growth on soil treated with digestate. Figure 3 below shows the plant at 35 days after emergence.

**Figure 3:**
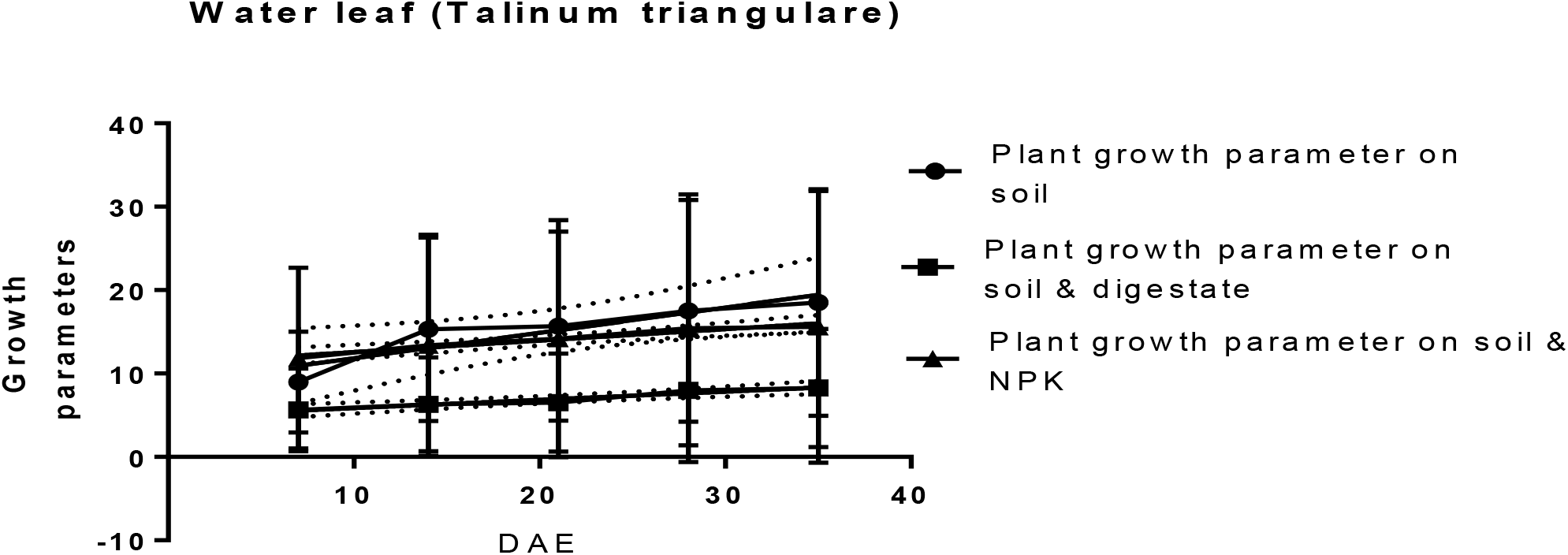
Water leaf (*Talinum triangulare*) at 35 Days of plant emergence

**Table 5:**
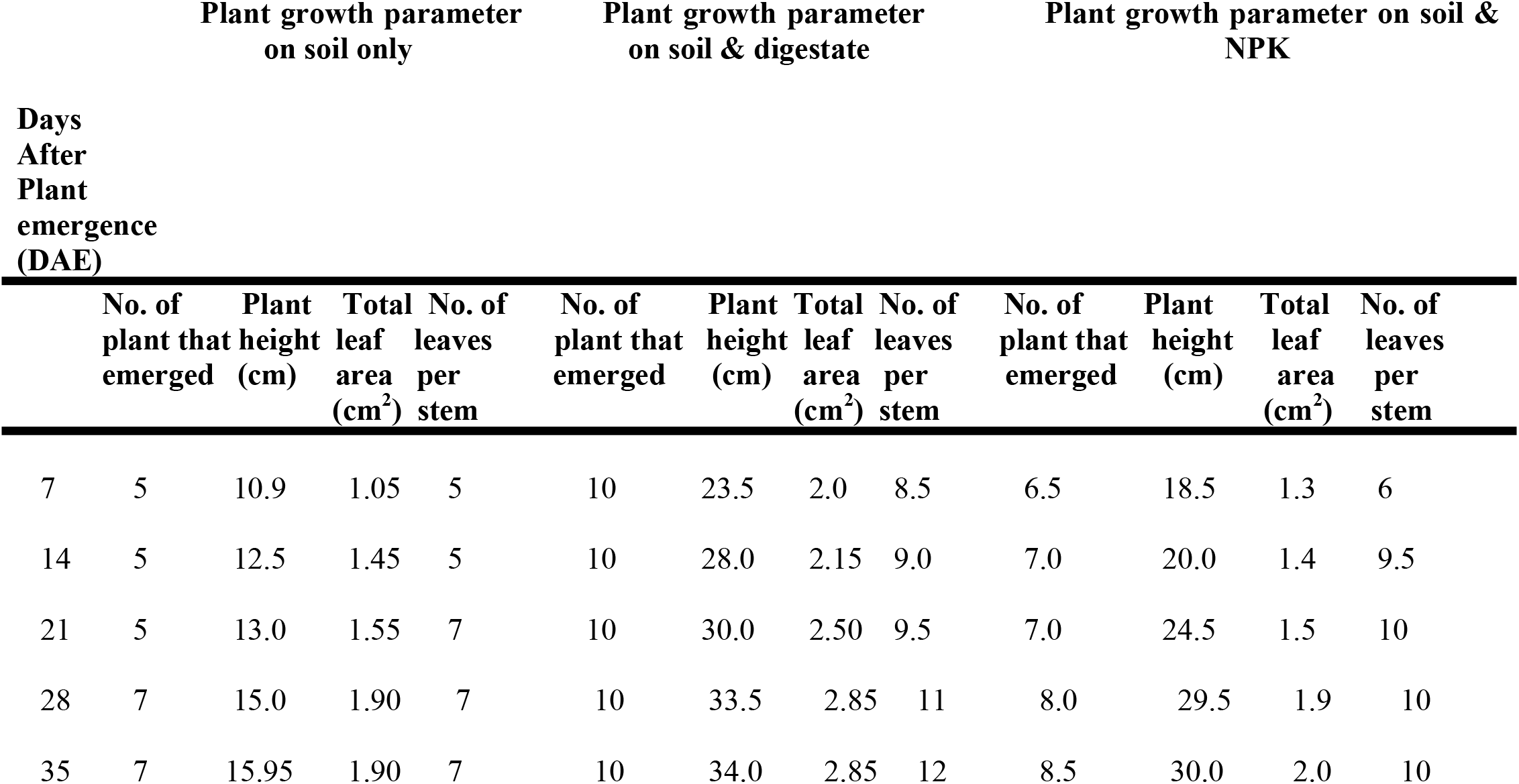
Tabular presentation of the growth parameter for water leaf (*Talinum triangulare*) on different treatments from day 7 to 35 DAE.

#### 3.2.3 Tabular presentation of Efo shoko (*Bot-celosia argentia*) growth parameter on different treatments

Table 6 below shows the growth parameters of Efo shoko (*Bot-celosia argentia*) on different treatments with a peak in the value of plant number on soil + digestate treatment (20) and soil+NPK (20) on the first week and gradually reduced to 16 & 10 respectively on the 35^th^ day. Other parameters such as number of leaves and plant height show equal value for all treatments. This plant had a stunted growth, at 14 DAE it couldn’t increase any further in height rather it reduced. Figure 4 below shows the plant at 35 days after plant emergence.

**Figure 4:**
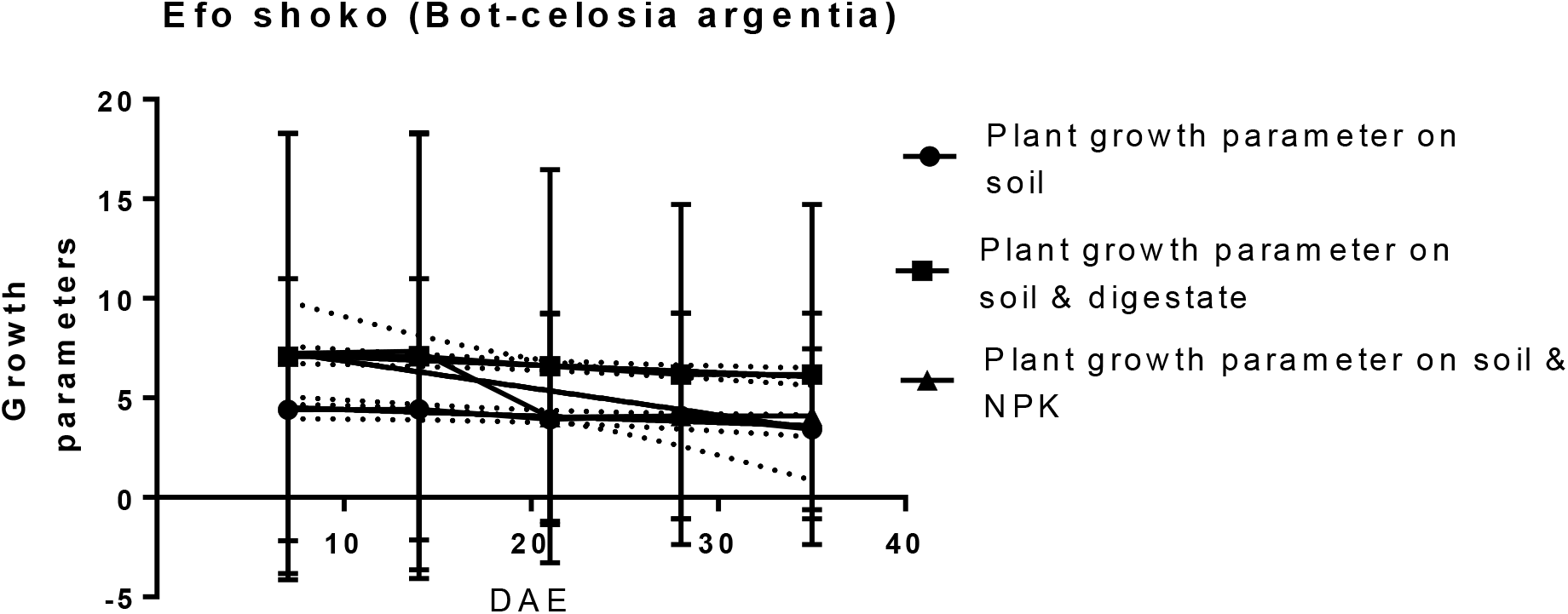
Efo Shoko (*Bot-celosia argentia*) at 35DAE

**Table 6:** Tabular presentation of the growth parameter for Efo shoko (*Bot-celosia argentia*) on different treatments from day 7 to 35 DAE.

| Days<br>After<br>Plant<br>emergence<br>(DAE) | Plant growth parameter<br>on soil only |  |  |  | Plant growth parameter<br>on soil & digestate |  |  |  | Plant growth parameter on soil &<br>NPK |  |  |  |
| --- | --- | --- | --- | --- | --- | --- | --- | --- | --- | --- | --- | --- |
|  | No. of<br>plant that<br>emerged | Plant<br>height<br>(cm) | Total<br>leaf<br>area<br>(cm<sup>2</sup>) | No. of<br>leaves<br>per<br>stem | No. of<br>plant that<br>emerged | Plant<br>height<br>(cm) | Total<br>leaf<br>area<br>(cm<sup>2</sup>) | No. of<br>leaves<br>per<br>stem | No. of<br>plant that<br>emerged | Plant<br>height<br>(cm) | Total<br>leaf<br>area<br>(cm<sup>2</sup>) | No. of<br>leaves<br>per<br>stem |
| 7 | 12 | 1.0 | 0.2 | 2 | 20 | 1 | 0.2 | 2 | 20 | 1.5 | 0.2 | 2 |
| 14 | 12 | 1.0 | 0.3 | 2 | 20 | 1 | 0.3 | 2 | 20 | 1.8 | 0.2 | 2 |
| 21 | 10 | 1.5 | 0.3 | 2 | 18 | 1.5 | 0.3 | 2 | 10 | 1.8 | 0.3 | 2 |
| 28 | 10 | 2.0 | 0.3 | 2 | 16 | 2 | 0.5 | 2 | 10 | 2.0 | 0.3 | 2 |
| 35 | 8 | 2.0 | 0.3 | 2 | 16 | 2 | 0.5 | 2 | 10 | 2.0 | 0.3 | 2 |

#### 3.2.4 Tabular presentation of Efo tete (Amaranthus hybridus) growth parameter on different treatments

Table 7 below is the tabular presentation of Efo tete (*Amaranthus hybridus*) plant growth parameter on different treatments obtained from day 7 to 35 DAE, with increase in the plant number of which soil + digestate treatment is the highest from 7 to 14 DAE. Other parameter such as the leaf number and leaf area for all treatments were very low and did not increase further after day 14. Figure 5 below shows the plant at 35 days after plant emergence.

**Figure 5:**
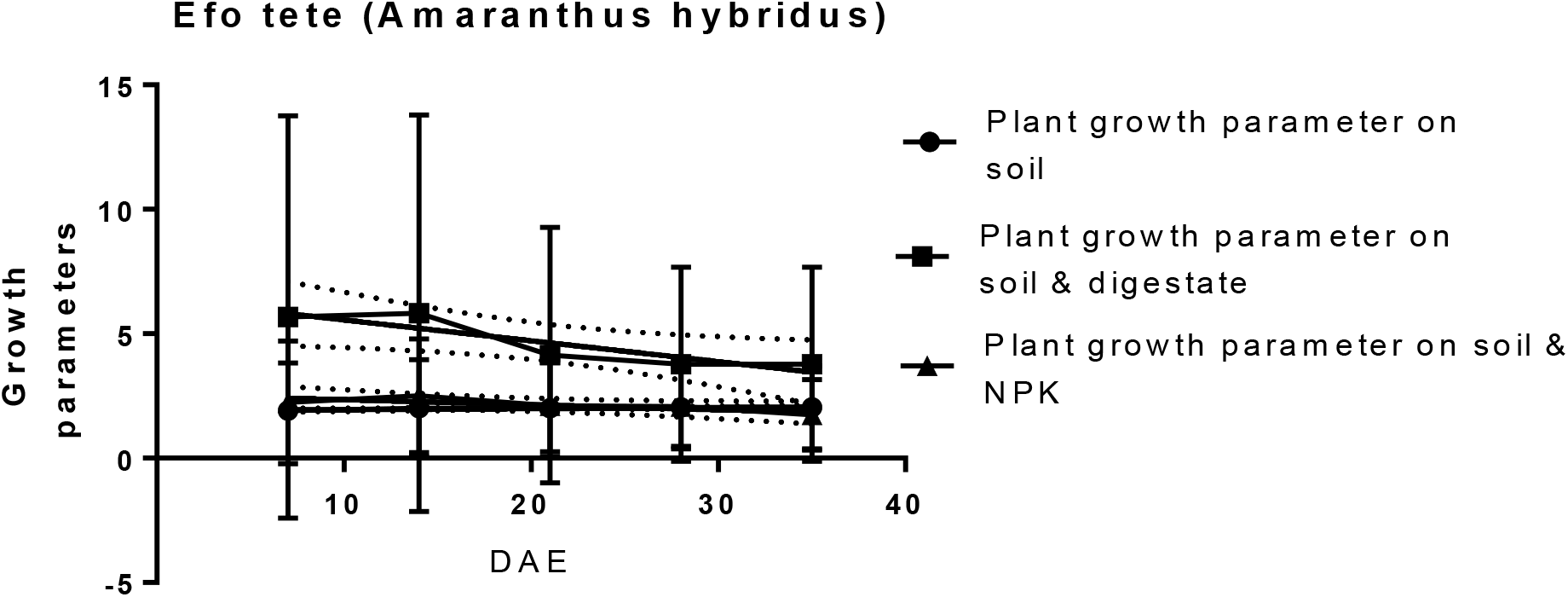
Efo Tete (*Amaranthus hybridus*) at 35DAE

**Table 7:** Tabular presentation of the growth parameter for Efo tete (*Amaranthus hybridus*) on different treatments from day 7 to 35 DAE.

| Days After Plant emergence (DAE) | Plant growth parameter on soil only |  |  |  | Plant growth parameter on soil & digestate |  |  |  | Plant growth parameter on soil & NPK |  |  |  |
| --- | --- | --- | --- | --- | --- | --- | --- | --- | --- | --- | --- | --- |
|  | No. of plant that emerged | Plant height (cm) | Total leaf area (cm<sup>2</sup>) | No. of leaves per stem | No. of plant that emerged | Plant height (cm) | Total leaf area (cm<sup>2</sup>) | No. of leaves per stem | No. of plant that emerged | Plant height (cm) | Total leaf area (cm<sup>2</sup>) | No. of leaves per stem |
| 7 | 4 | 1.5 | 0.2 | 2 | 15 | 1.5 | 0.5 | 2 | 5 | 1.5 | 0.5 | 2 |
| 14 | 4 | 1.9 | 0.1 | 2 | 15 | 2.0 | 0.5 | 2 | 5 | 2.0 | 0.5 | 2 |
| 21 | 4 | 1.9 | 0.1 | 2 | 10 | 2.0 | 0.4 | 2 | 4 | 2.0 | 0.3 | 2 |
| 28 | 3 | 3.0 | 0.1 | 2 | 8 | 3.0 | 0.3 | 2 | 3 | 3.0 | 0.2 | 2 |
| 35 | 3 | 3.0 | 0.1 | 2 | 8 | 3.0 | 0.3 | 2 | 2 | 3.0 | 0.2 |  |

### 3.3 Effect of different treatments on the growth of water leaf

From the table 8 below, it can be seen that there is a significant difference between the treatments of soil+digestate, soil+NPK and soil without treatment used for the growth of water leaf plant from day 7 to day 35 at p≤0.05.

**Table 8:**
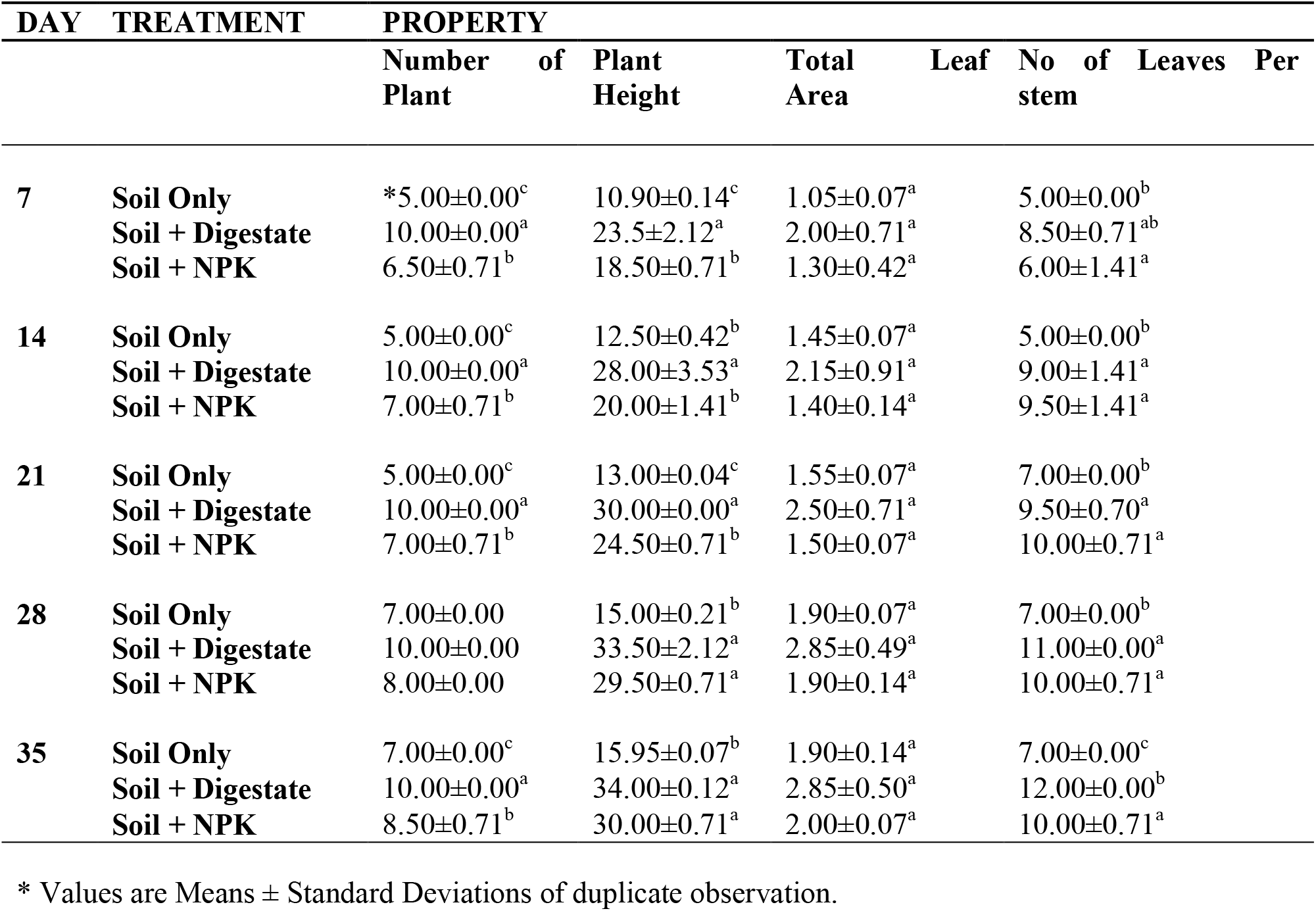
Effect of different treatments on the growth of water leaf.

*Means with same superscript (a, b, c) letter down each column are not significantly different from each other at p≤0.05 using New Duncan Multiple Range Test.*

### 3.4 Bacterial community in the digestate

#### 3.4.1 Taxonomic distribution of bacterial community in the digestate at phylum level

It can be inferred that *Firmicutes* and *Proteobacteria* are the most abundant, with *Bacteriodetes* (2%) been the least abundant. Detail on the taxonomic distribution at phyla level is summarized below in figure 6.

**Figure 6:**
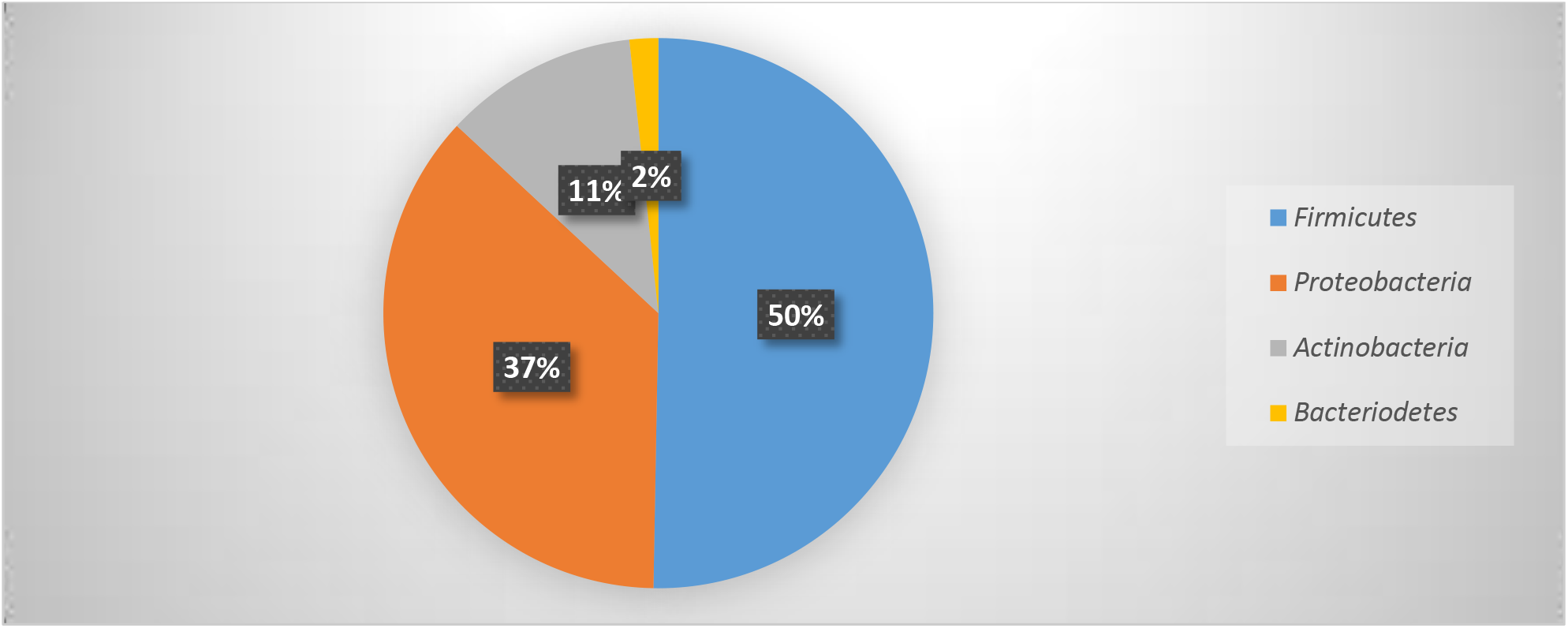
Pie chart showing the relative abundance of the top five (5) phyla within the bacterial community in the digestate.

#### 3.4.2 Taxonomic distribution of bacterial community in the digestate at class level

From the figure 7 below, it can be inferred that *Clostridia* (27%) and *Bacilli* (26%) are most abundantly present followed by *Actinobacteria*. The class *Bacteriodia* happen to be the least in abundance with (7%).

**Figure 7:**
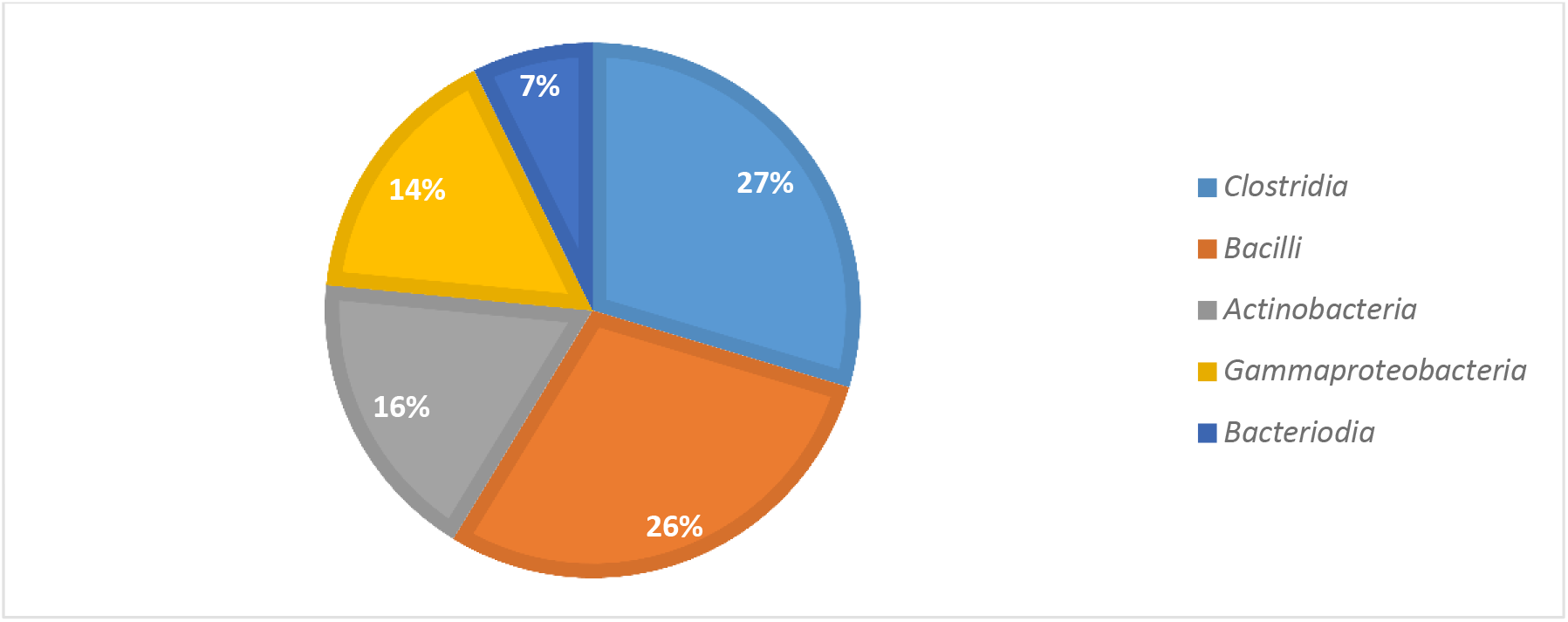
Pie chart showing the relative abundance of the top five (5) classes within the bacterial community in the digestate.

## DUSCUSSION

Digestate produced from biogas processing using readily sourced organic waste such as pineapple peel, food waste, sawdust and animal waste (cow dung) has been used as a biofertilizer for selected vegetables. This study is important in determining how effective a digestate produced from co-digested biomass is as a biofertilizer for vegetable farming. Most of the research carried out using digestate derived from food waste and animal waste was mainly used for cereal farming (Zhang, 2017; Katarzna *et al*., 2018) with few studies on vegetables (Losak *et al*., 2016). The selected vegetables used for the pot test were Ewedu (*Corchorus olitorius*), Efo Shoko (*Bot-celosia argentia*), Efo tete (*Amaranthus hybridus*) and Water leaf (*Talinum triangulare*). The effectiveness of the digestate in the growth of these vegetables was compared to that of inorganic chemical fertilizers.

From the nutritional study carried out, it can be shown that the values of phosphorus (p) and potassium (k) were substantially higher in soil (50.65mg/100g and 67.41ppm, respectively) than in digestate 18.44mg/100 g for phosphorus and 55.31ppm for potassium, which decreased to 16.11mg/100 g for phosphorus and 42.67ppm for potassium in soil and digestate mixtures. Nitrogen is needed by the plant for optimum growth, being the major component of amino acids is an essential nutrient for photosynthesis in the plant (Simon *et al*., 2014). Nitrogen content was found to be 1.62% in soil and digestate mixtures, which is an elevated value relative to nitrogen content in soil alone (0.94%). The pH of the digestate is based on the type of biomass used. In this analysis, saw dust, cow dung and food waste were used and the pH of the resulting digestate (8.1) is consistent with the previous Pognani *et al*. (2009) research. The nutrient composition of the digestate obtained in this study shows the presence of three major nutrients (Nitrogen, Potassium and Phosphorus). This suggests that digestate is rich in nutrients and has the potential to increase soil microbial and nutrient status when used as fertilizer, especially in nutrient-depleted soils. Several studies have identified anaerobic digestate potentials as an effective substitute for inorganic chemical fertilizers known to have adverse effects on the environment (Westphal *et al*., 2016).

The Phyto assessment of the four selected vegetables used in this study compared growth parameters between plants grown in different soil treatments, including soil fertilized with digestate, chemical fertilizer (positive control) and soil without fertilizer (negative control). The performance of the plants in the different treatments can be measured by their growth parameters. Waterleaf (*Talinum triangulare*) plant as shown in Figure 3 had a better performance on soil treated with digestate compared to its performance on soil only and soil + NPK. This is analogous to the pot test experiment of the Lettuce plant using digestate as a biofertilizer by Trinchera *et al*. (2013). The lettuce plant grown on soil treated with digestate grew stronger than the urea-treated soil plant (chemical fertilizer used as a positive control). The Duncan multiple range test was used to statistically evaluate the water leaf (*Talinum traingulare*) data to see a significant difference between the treatments used. The findings showed a significant difference at (p≤0.05) suggesting a comparison between the use of soil+digestate and soil+NPK as a treatment for the growth of water leaf. On the other hand, the Ewedu plant (*Corchorus olitorius*) performed poorly on soil treated with digestate as seen from the tabular presentation of the growth parameter in Table 4, compared to the chemical fertilizer (positive control) and the unfertilized soil (negative control). This may be the effect of the digestate on the soil type, as the digestate is regarded as a complex material with a beneficial effect on the physical, chemical and biological properties of the soil, depending on the soil type (Makadi *et al*., 2008). However, both Efo shoko (*Bot-celosia argentia*) and Efo tete (*Amaranthus hybridus*) had a stunted development in both treatments as seen from the tabular presentation of their growth parameters in Tables 6 and 7. This could be due to untreated digestate (Drosg *et al*., 2015) or over application of the digestate that inhibited the growth of certain vegetable forms, such as Efo tete (*Amaranthus hybridus*) and Efo shoko (*Bot-celosia argentia*), as it has been stated that caution must be taken in the application of digestate because it contains a high nutrient content and may lead to phytotoxicity over use (Nkoa, 2014; Insam *et al*., 2015). This is corroborated by earlier reports that anaerobic digestate contains a relatively high proportion of mineral nutrients, which gives the digestate a strong fertilizing capacity to replace inorganic fertilizers, especially due to nutrient retention even after raw material digestion (Alburquerque *et al*., 2012). The use of digestate as a biofertilizer for these vegetables is consistent with the Suarez *et al*. (2015) study, that the application of biofertilizer stimulates plant growth by various mechanisms such as atmospheric nitrogen fixation, phosphorous solubilization and mobilization, iron sequestration by siderophores and phytohormone production. Over-reliance on inorganic fertilizers, particularly in the tropics that are on the rise, has resulted in a decline in soil fertility, environmental degradation and even a contribution to the release of greenhouse gases (Chandini *et al*., 2019). Organic fertilizers, such as biofertilizers with biofertilizers potential, are therefore necessary for adequate supply of nutrients to plants and also function as an amendment to increase soil humus, coupled with its eco-friendliness compared to inorganic fertilizers (Sun *et al*., 2015).

Metagenomic study of the digestate revealed the presence of Bacteria, with phylum *Firmicutes* in abundance (50%) followed by *Proteobacteria* (37%), *Actinobacteria* (11%) and *Bacteriodetes* been least in abundance (2%). Abundant members of the class include *Clostridia* (27%) and *Bacilli* (26%). This is followed by a study by Treu *et al*. (2016) which, among others, had *Firmicutes* and *Clostridia* as the most prevalent phylogenetic class. *Firmicutes* were the most abundant phyla in the digestate and are most influential in the biogas reactor, which is consistent with previous research by Krober *et al*. (2009) and can be attributed to their ability to degrade polysaccharides and oligosaccharides. Hydrolysis is the first step towards the effective conversion of plant biomass to intermediate compounds which are further degraded to methane. It is well known that cellulose degradation (the most abundant plant polysaccharide) is performed in various habitats by species associated with phyla *Firmicutes* and *Fibrobacteres*. These species encode the cellulosome, one of the most widely recognized cellulase used for both adhesion and enzymatic purposes (Treu *et al*., 2016). Bacteria belonging to the phylum *Proteobacteria* which are the second in abundance have *Gammaproteobacteria* (14%) as the fourth abundant class and are said to use glucose, acetate and propionate during the AD phase, while the majority of the bacteria belonging to the phyla *Bacteriodetes* which is the least in abundance (2%) are known to produce lytic enzymes and acetic acid. According to De Francisci *et al*. (2015), the central function of *Bacteroidetes*, together with *Firmicutes*, confirms the results of previous studies which indicated that microbes belonging to these phyla contribute to the decomposition of cattle manure, consisting mainly of plant biomass residues, in biogas reactors. *Clostridia* (27%) and *Bacilli* (26%) are the most abundant in the class level, followed by *Actinobacteria* (16%) and *Gammaproteobacteria* (14%) with *Bacteriodia* being the least abundant (7%).

The presence of these bacteria in the digestate correlates with the analysis of the microbial community in the anaerobic digestion of cow dung and mixed food waste by Roopnarain *et al*. (2019) although they recorded the abundance of *Clostridia* and *Bacteriodia. Staphylococcus* is the abundant genus (39%) with *Epidermitis* (0.99%) as the least abundant species followed by *Legionella* (11%) and *Micrococcus* the third abundant genus (8%) has the most abundant species *Luteus* (8.25%), other genus present also includes *Acinetobacter* (5%) which has the second abundant species *iwoffii* (1.36%) and *Pseudomonas* been the least abundant genus (4%) with the least abundant species *stutzeri* (0.57%). The existence of these bacteria suggests that the digestate is rich in microorganisms used as a microbial inoculant for soil fertilization and nutrient enhancement. Suitable inoculants such as *Bacillus, Clostridium,* and *Pseudomonas* have been found in the digestate and these bacteria are known to accelerate the microbial process in the soil by increasing the supply of nutrients that can be assimilated by plants (TNAU, 2008). *Clostridium* species are free nitrogen fixings, while *Bacillus* species are phosphate solubilizers (Alfa *et al*., 2014). In the meantime, the presence of *Staphylococcus*, while having the least species (*epidermidis*), appears to be a possible risk of digestate when used in food crops, thus requiring the treatment of digestate or deactivation of *Staphylococcus* in the AD system, as reported by Kirby *et al*. (2019) before use, so as not to have it in food and pose a health risk to the end user.

## Conclusion

The digestate produced from the different waste biomass used in this study has shown from the results that the digestate is rich in mineral elemental composition, such as Nitrogen, Phosphorus and Potassium, as well as a favorable pH for plant growth, and contains beneficial soil bacteria such as Nitrogen fixers and Phosphate solubilizers. All of these make the digestate a possible biofertilizer. While other vegetables may not survive on soil treated with digestate, which may be due to the untreated digestate used or the over use of digestate, which may have a toxic effect on some of the vegetables, resulting to their stunted growth. Further treatment of the digestate to remove pathogens such as *staphylococcus* as revealed in the methagenomic analysis and given a more controlled application management condition, these vegetables may thrive and possibly perform better than when grown on soil with chemical fertilizer just as the waterleaf (*Talinum triangulare*). The existence of beneficial bacteria and nutrients suggests that digestate is a biofertilizer, while the existence of *staphylococcus* involved in digestate is a major health concern, it is therefore suggested that more research on the treatment of digestate should be performed to understand how feasible it will be to treat digestate from *staphylococcus* without causing possible damage to other beneficial bacteria, or during application of the digestate it should be in such a way that it will not be spread directly on the vegetables. Consequently, given the significant difference in the treatment of digestate and chemical fertilizers used in this report, it is concluded that the use of digestate as a biofertilizer would minimize the over-dependence of inorganic chemical fertilizers in Nigeria and other countries in the tropics, thus overcoming the challenges of environmental contamination and reducing soil quality.

## Author’s contribution

All authors were fully involved in bringing out the authenticity of this study, take responsibility for the accuracy and integrity of the data/data analysis. F.C. Raymond worked on the conceptuality, sourcing of data, analyse data, develop theories, performed the investigation, funded and wrote the manuscript. O.M. Buraimoh supervised the research study, worked on the conceptuality, develop theories, verified the research method, help analyse data, encouraged F.C. Raymond and O.S. Akerele to source current method used to carry out this research and edited the manuscript. O.S. Akerele help develop theories, supported in the investigation, helped F.C. Raymond to source and analyse data and proof read the manuscript. O.S. Illori verified the research method, help to supervise the research and proof read the manuscript. O.T. Ogunipe help to supervise the research, verify the research method and help in editing the manuscript.

## Acknowledgement

This research was self-funded, and thanks to Sir Raymond Ferdinand for the financial support.

## APPENDIXES

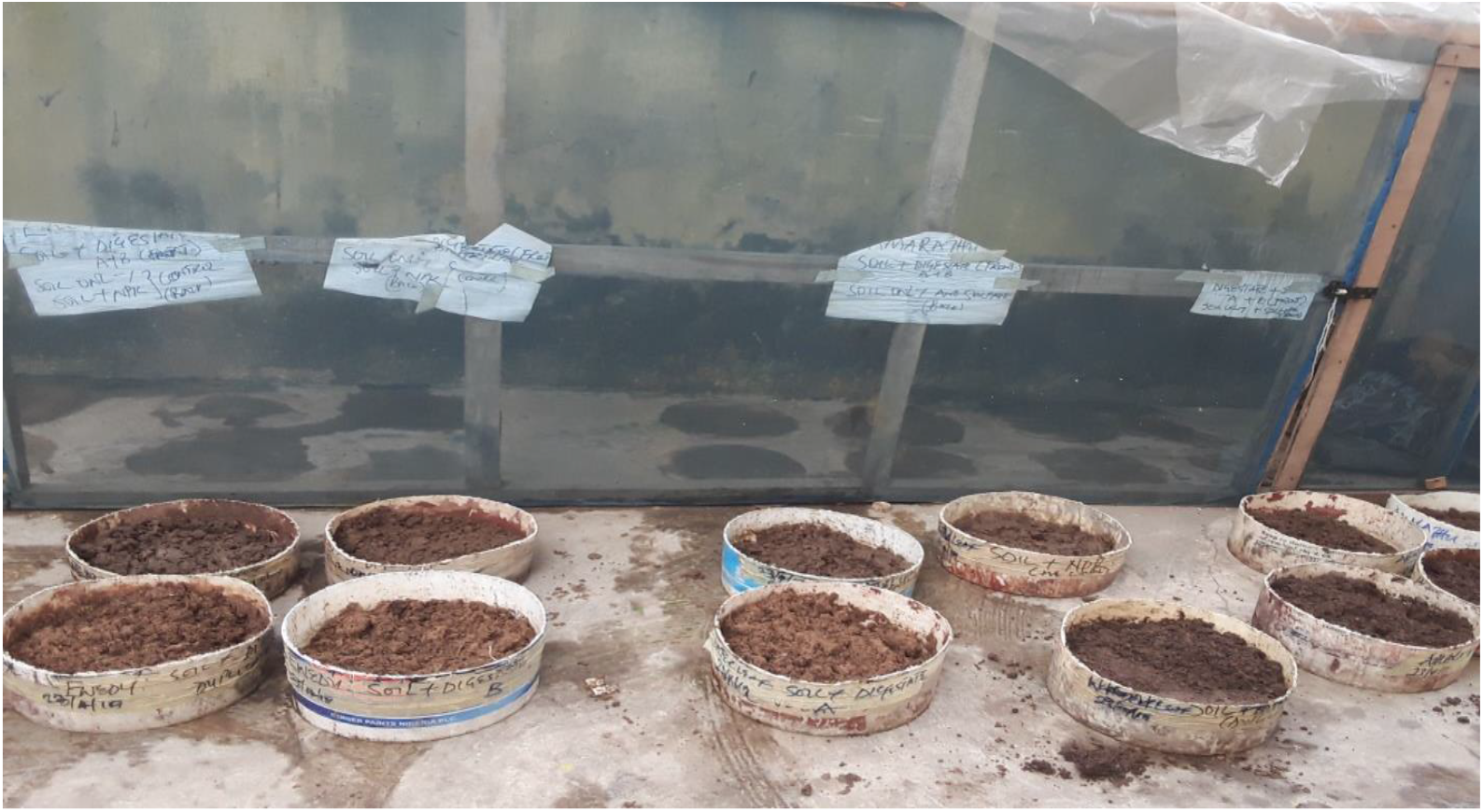

Ewedu (*Corchorus olitorius*) growth parameter

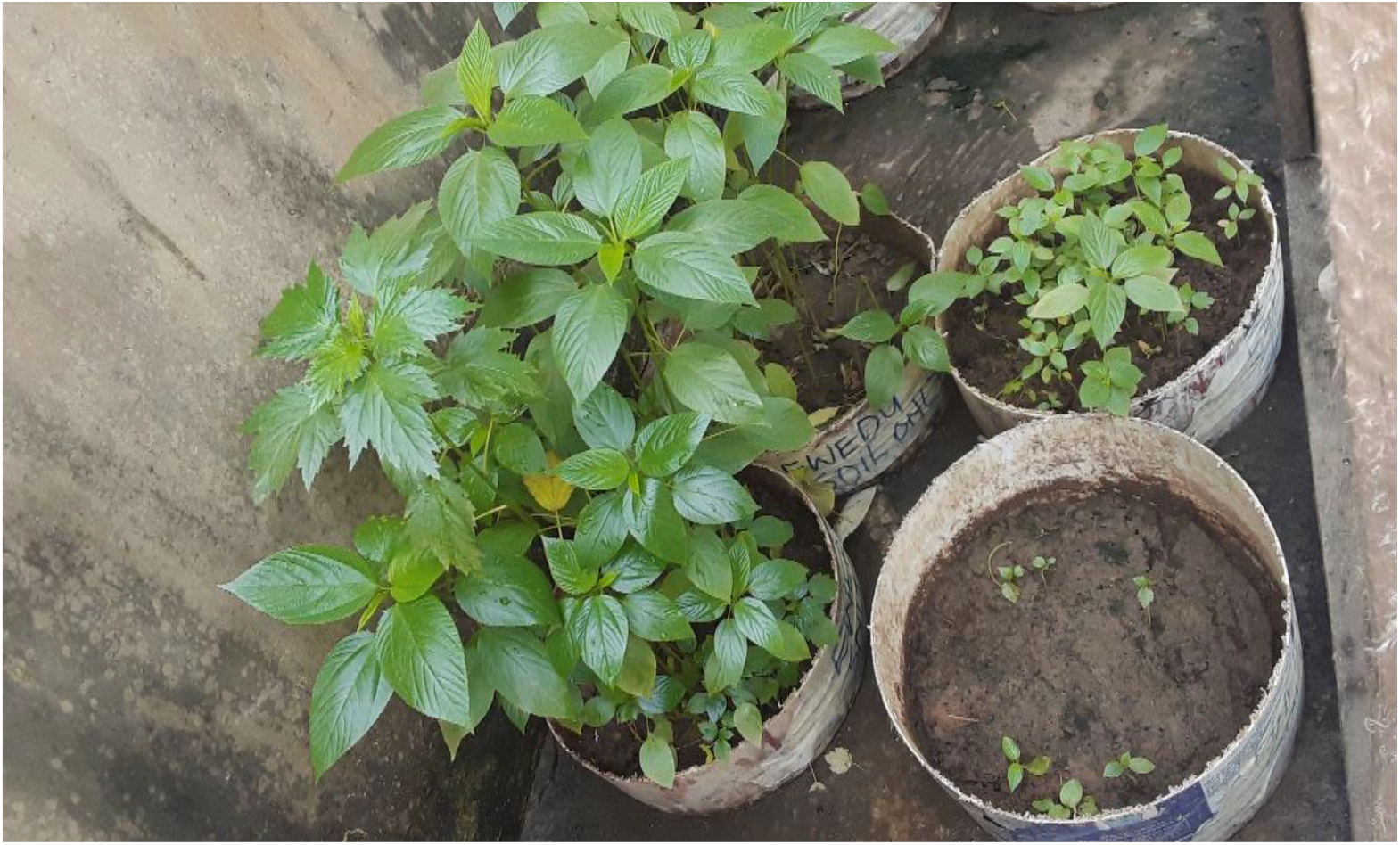

Water leaf (*Talinum triangulare*) growth parameter

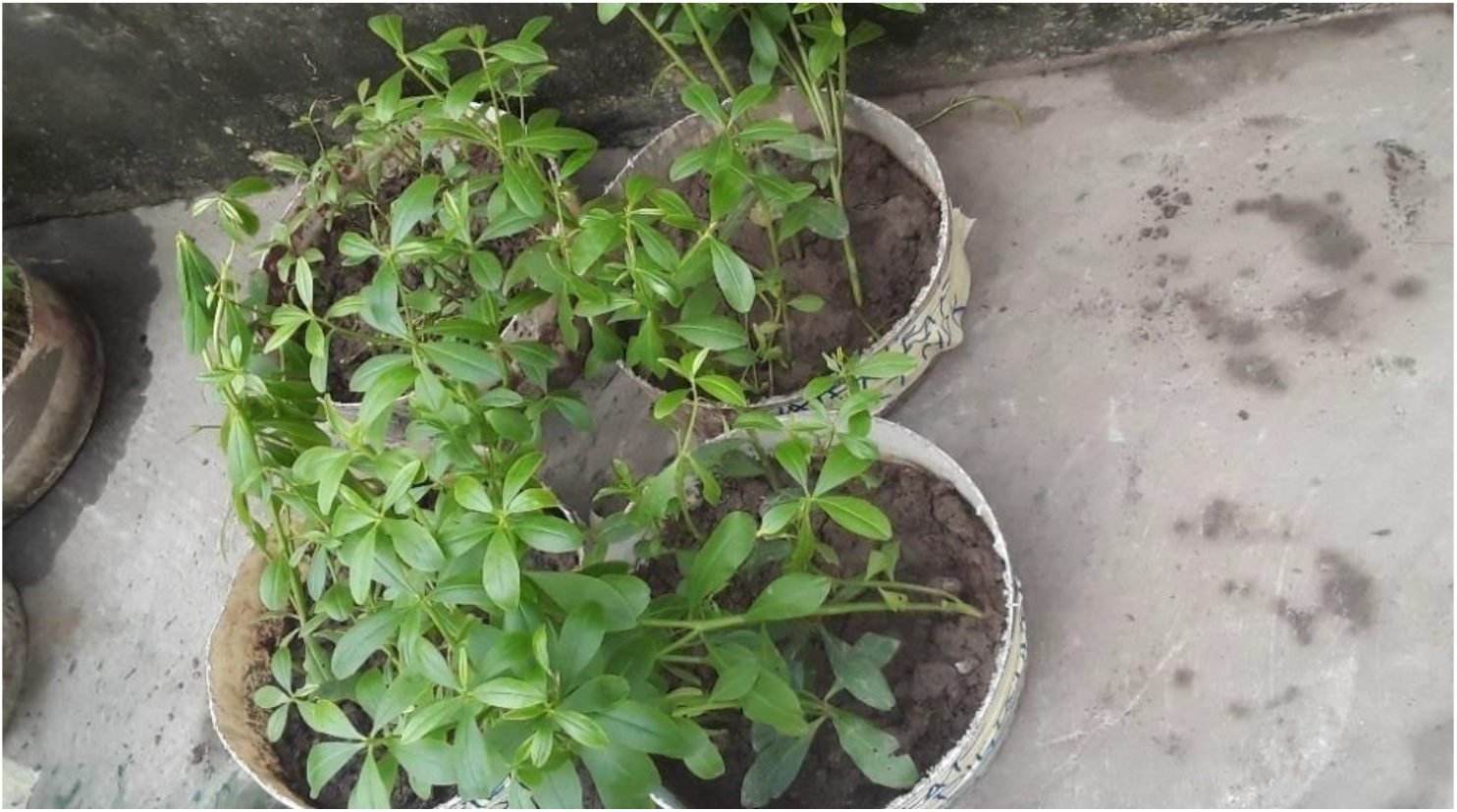

Efo shoko (*Bot-celosia argentia*) growth parameter

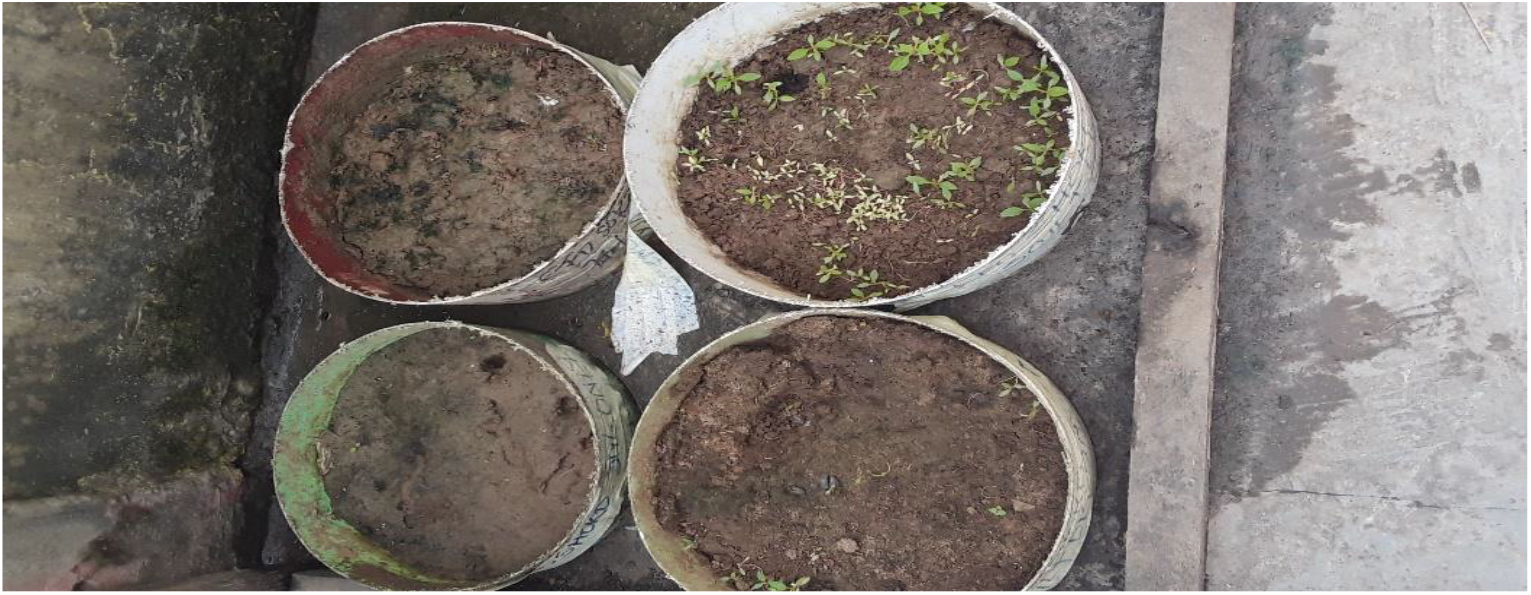

Efo tete (*Amaranthus hybridus*) growth parameter

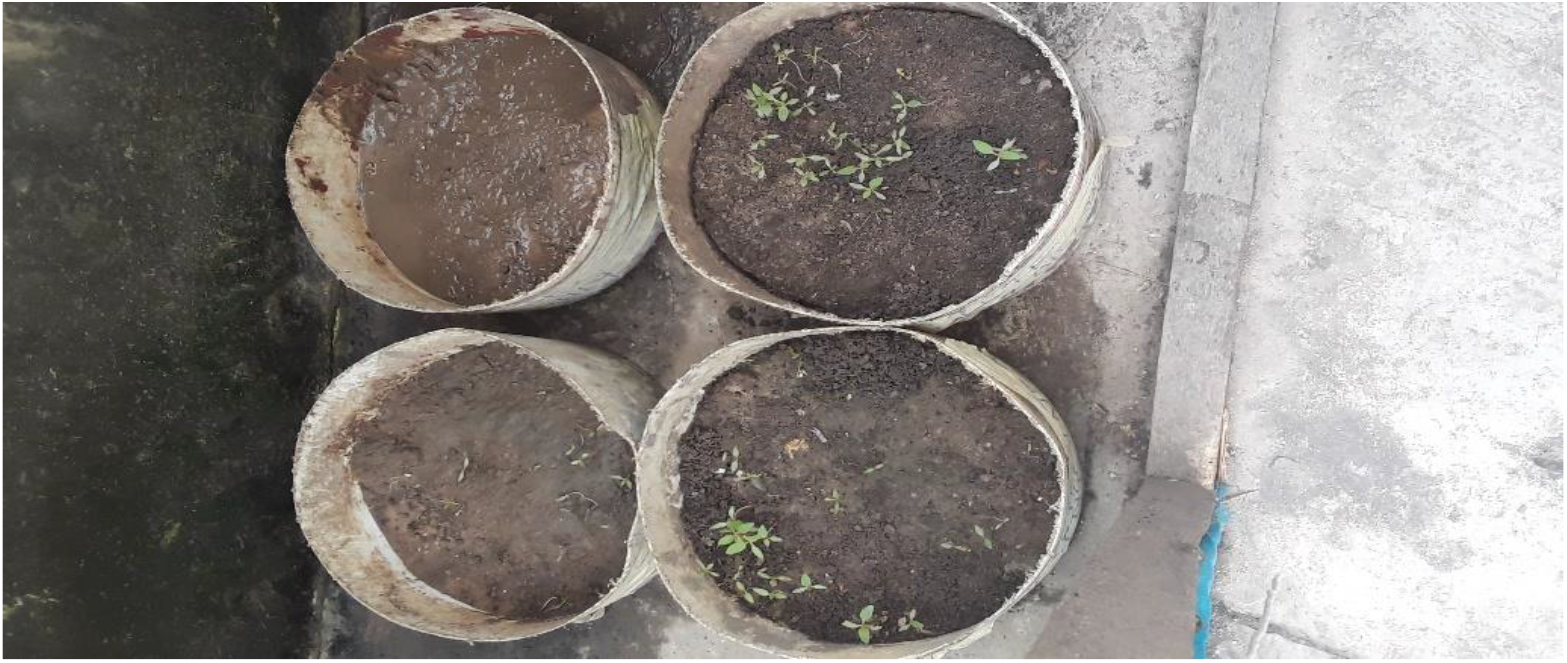

